# Bone remodeling: A tissue-level process emerging from cell-level molecular algorithms

**DOI:** 10.1101/318931

**Authors:** Clemente F. Arias, Miguel A. Herrero, Luis F. Echeverri, Gerardo E. Oleaga, José M. López

## Abstract

Human skeleton undergoes constant remodeling during the whole life. By means of such process, which occurs at a microscopic scale, worn out bone is replaced by new, fully functional one. Multiple bone remodeling events occur simultaneously and independently throughout the body, so the whole skeleton is completely renewed about every ten years.

Bone remodeling is performed by groups of cells called Bone Multicellular Units (BMU). BMUs consist of different cell types; some are specialized in destroying old bone, whereas others produce new bone to replace the former. The whole process is tightly regulated so that the amount of new bone produced exactly balances that of old one removed and bone microscopic structure is maintained.

To date, many regulatory molecules involved in bone remodeling have been identified, but the precise mechanism of BMU operation remains to be fully elucidated. Given the complexity of the signaling pathways already known, the question arises of ascertaining whether such complexity is an inherent requirement of the process, or a consequence of operational redundancy.

In this work we propose a minimal model of BMU function which involves a small number of signals and accounts for fully functional BMU operation. Our main assumptions are i) at any given time, any cell within a BMU can select only one among a reduced choice of decisions: divide, die, migrate or differentiate, ii) such decision is irreversibly determined by depletion of an appropriate internal inhibitor and iii) the dynamics of any such inhibitor is coupled to that of a few external mediators. It is shown that efficient BMU operation then unfolds as an emergent property, which results from individual decisions taken by cells in the BMU unit in the absence of any external planning.

**Author summary:** Our skeleton is a living organ that is being renewed throughout our life. This task is accomplished by teams of bone cells termed as Bone Multicellular Units (BMUs) that are recruited when and where needed, to operate at places where bone has lost functionality either for an excess of mechanical stress or because loss of activity. Once assembled, BMU remove old bone and replace it by new one, and disband as soon as their mission has been accomplished. No single bone evades BMU screening, so that the whole human skeleton is completely renewed approximately every ten years.

It is natural to wonder how such robust and fascinating process is regulated. Many signaling pathways involved in bone remodeling have been identified so far, but whether all of them are necessary for BMU operation remains unclear. In this work we show that just a reduced number of such signals could suffice for that purpose. This suggests that a large degree of redundancy might have been kept in place, perhaps as a consequence of different convergent strategies developed in the course of evolution.

## Introduction

The human skeleton is a complex structure made up of 206 bones, which provides a supporting framework for the body. It acts as a shield to protect internal organs and plays a crucial role in locomotion by mediating the force arising from muscle contraction. In spite of its inert appearance, bone is an extremely dynamic tissue that is continuously being remodelled to adapt to changing mechanical demands. Such remodeling, which is carried out at a microscopic scale, consists in the removal of low-performing bone and its replacement by new, fully functional one. This task is fulfilled by suitable agents called for that purpose, as described below.

Bone tissue is formed by a mineralized matrix that has been hardened to provide a supporting function. There are three key cell types that are responsible for matrix production, maintenance and remodeling: osteoclasts, osteoblasts and osteocytes which perform different homeostatic functions [1–3]. Osteoclasts, recruited when needed from their cell precursors, are in charge of degrading dysfunctional bone, whereas synthesis of new bone to replace the former is carried out by osteoblasts. Osteocytes, the most abundant bone cells, form a three-dimensional interconnected network throughout the bones. They act as mechanosensors that monitor mechanical stress within bone tissues, and react to changes in both the amount and the direction of loading applied on bones.

A key event that triggers bone remodeling is osteocyte cell death (apoptosis) occurring over comparatively small length scales and resulting, for instance, from unusual mechanical load or from micro-fractures induced by physical exercise. Concerning the first situation, it should be mentioned that the relation between osteocyte apoptosis and load applied is known to be U-shaped. This means that mechanical stresses within a normal physiological range prevent it, whereas those above or below this range induce it [4–6]. In traumatic bone fractures, a considerable amount of osteocytes are eliminated and alert signals are produced that recruit immune cells and result in an inflammatory response. In such case, an alternative mechanism of bone formation is triggered, where other cell types are implicated [7]. We shall not deal with this case here, since we will be concerned with homeostatic bone remodeling at smaller scales. The manner in which such process occurs is now succinctly described.

Following osteocyte apoptosis in a microscopic region (termed as Bone Remodeling Compartment (BRC), which is often about 400 microns wide), organic teams called Bone Multicellular Units (BMU) are locally recruited [8, 9]. Each BMU consists of several morphologically and functionally different cell types, mainly osteocytes, osteoblasts and osteoclasts, that act in coordination over the BRC to replace old bone by new one [10, 11]. Not all cell types required are initially at place. In fact, prior to remodeling a normal presence of osteocytes produces an inhibitory effect that keeps osteoblasts deactivated and prevents osteoclast precursors, which eventually will give raise to acting osteoclasts, from being called in. However, osteocytes apoptosis over the BRC results in a drop of such inhibitory action, which has several consequences. To begin with, it leads to osteoblasts activation and to the recruitment of osteoclasts precursors arising from the bone marrow. Such precursors subsequently differentiate into mature osteoclasts, which start the erosion (resorption) of adjacent bone, leading to the appearance of the so-called cutting cones [12]. In a later stage, activated osteoblasts, marching in the wake of bone-destroying cutting cones, will replenish the cavity left behind the latter by secreting an osteoid matrix, a precursor of new bone. Some of the active osteoblasts become trapped in the matrix that they secrete and eventually differentiate into osteocytes [13], which in turn will be instrumental in the subsequent mineralization of the matrix surrounding them, thus concluding a local bone remodeling event. The whole process is summarized in Figure 1 below

**Fig 1.**
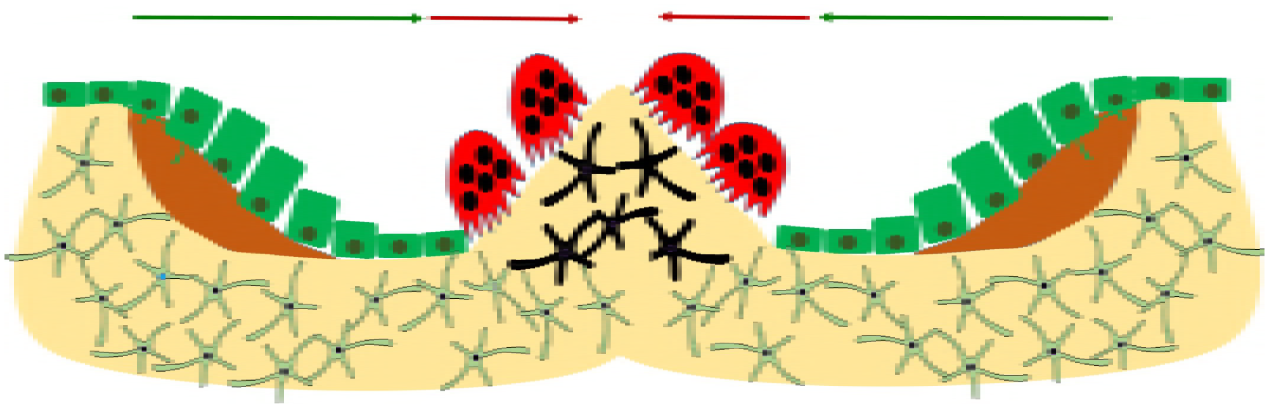
A sketch of BMU operation after a group of osteocytes has undergone apoptosis near bone surface. Bone resorption by osteoclasts (red arrow) and bone formation by osteoblasts (green arrow) are spatially and temporally coupled. Bone remodeling is initiated when osteoclast precursor cells are recruited to the altered bone surface (black stellate cells) and fuse to form mature, bone resorbing osteoclasts (red cells) that attach to the surface. Mature osteoclasts degrade the mineralized matrix and produce resorption pits also called resorption bays or Howship’s lacunae. Once osteoclasts have degraded the target area, they undergo apoptosis, and osteoblasts situated behind them first secrete osteoid matrix and subsequently differentiate into osteocytes

It should be stressed that the process just recalled is exquisitely regulated. It only takes place where needed and the amount of new bone generated exactly balances that of old bone destroyed, thus warranting that no net changes in bone mass nor mechanical stress remain after each remodeling cycle. Any imbalance between bone resorption and bone formation might lead to pathological disorders. For instance, five important human diseases: osteoporosis, renal osteodystrophy, Paget’s disease, osteopetrosis and rickets are associated to malfunctions in bone remodeling.

To date, and in spite of substantial progress achieved during the last decades, the mechanisms of cell signaling that regulate BMU operation remain only partially understood. It is known that such regulation is of a local nature, since remodeling is simultaneously and independently occurring at different places, so that the whole adult skeleton is renewed approximately every ten years [14]. The local nature of that process suggests that it should be a consequence of individual cell decisions that must be somehow coordinated at a population level. For instance, osteoblasts should not start secreting osteoid matrix before osteoclasts destruction task is finished. This raises at once the questions of how such coordination can be achieved, and what internal circuitry within a BMU keeps such unit operative when remodeling is needed, and shuts it off immediately afterwards. We shall address these issues in the following sections.

Specifically, in this work we propose and analyse a simple, space-dependent model that is able to reproduce BMU operation and requires only of a small number of signaling cues. In our model, BMU function is shown to result from a limited number of individual decisions (divide, die, migrate or differentiate) taken by any of the cells involved as a consequence of the coupled dynamics of external cues and internal decision inhibitors. Simplicity is a key goal here. We are in fact interested in describing a minimal software sufficient to ensure BMU operation and involving generic molecules able to elicit actions that have already been experimentally observed. We therefore not aim at providing a comprehensive model of bone remodeling where most (if not all) possibly involved pathways are retained. Rather, we are intent to identify the minimal circuitry needed to keep a BMU operative. We believe that this may be a useful complement to previous work (see for instance [15–17] and references therein) where attention is paid to building models that include as many biological ingredients as possible. We hope that a combined use of both approaches (comprehensive and minimality-driven) could shed further light into our understanding of bone remodeling and on the control of this process when such action may be needed.

## Models

### The role of molecular signaling in bone remodeling

To date, a considerable number of signals involved in BMU operation have been identified in the literature [10, 14, 18, 19]. For instance, sclerostin is secreted by normal osteocytes and is known to inhibit both osteoblast activation and osteoclast recruitment from precursor cells in the bone marrow [10, 20]. This inhibitory effect prevents the activation of BMUs in regions where bone remodeling is not needed. In fact, only when an appropriate number of osteocytes undergo apoptosis in a given area, due for instance to micro-fractures or changes in mechanical load, will the inhibitory effect of sclerostin be significantly depleted, thus entailing the activation of a BMU and the subsequent initiation of a bone remodeling event.

On the other hand, a set of signals known under the generic name of Bone Morphogenetic Proteins (BMPs) have been shown to play a key role during early stages of bone remodeling. Some members of this family, such as TGF-*β* and TGF-*α*, are known to foster differentiation of mesenchymal stem cells to osteoblasts [21, 22]. TGF-*β* can also induce migration of osteoblasts to the sites of bone formation during remodeling [23, 24] and inhibits osteoblast apoptosis [5]. TGF-*β* is mainly produced by osteocytes [20, 25], but it is also present in the bone matrix [21] and in platelets [26]. This last factor, together with other signals such as HMGB-1 [27] provides a link with inflammatory processes occurring at early stages of large fractures repair. Cytokines such as IGF-1, released from bone matrix, seem to play a similar role in activating osteoblast differentiation [28] and are necessary for their survival in vitro [29].

The next stage in the process of bone remodeling consists in the recruitment of osteoclasts. The main signals involved in this step are M-CSF and RANKL [19, 30, 31], that promote the differentiation of osteoclast precursors and the survival of activated osteoclasts. Since RANKL is mostly produced by activated osteoblasts [10, 30], osteoclasts will only be recruited to sites were bone remodeling has already been triggered. Besides inhibiting osteoblast activation, sclerostin also induces apoptosis of active osteoclasts [10, 20]. It may thus act as a signal that stops bone resorption when the cutting edge has reached the required depth in each remodeling event. Finally, osteoid matrix production by active osteoblasts, as well as differentiation of osteoblasts into osteocytes seem to be cell-density dependent [32], and have been suggested to be mediated by connexin, a molecule that circulates through gap junction communications between osteoblasts, as well as by sclerostin [18, 33]. Finally, various chemoattractants/chemorepulsors that drive osteoclasts away from the region where their precursors are recruited have been described in the literature (see for instance [34]). We shall use one such generic signal as part of our algorithm below.

Concerning signaling effects, one also has to bear in mind that: I) different signals can have redundant effects; for example both TGF-*β* and IGF-1 induce osteoblast differentiation [35], II) a given signal can have different effects on different cell types. For instance TGF-*β*, FGF and PDGF activate osteoblast and inhibit osteoclast action [26] and G-CSF is known to induce apoptosis and to inhibit differentiation in osteoblasts [31], and III) signals are not always provided by chemicals. In this context we have already remarked that osteocytes act as sensors that respond to changes in mechanical stress in bone [36, 37].

The list just provided of molecular mediators and their effects on BMU cell types, which is summarized in Table 1 below, is far from being complete. Moreover, knowing the identity of such mediators or describing their effects in qualitative terms is not enough to explain BMU operation during bone remodeling. In order to understand how a coherent collective plan of action emerges at a multicellular scale, quantitative aspects of the process need to be taken into account. Indeed, for any given set of signals involved, the amount of bone to be resorpted and produced in different remodeling events can be highly variable [10]. This implies in turn that the number and activity of cells recruited in a BMU should change to suit the needs at each particular remodeling process [25, 26, 38, 39].

**Table 1.** A brief summary of the main signals involved in BMU operation as described in the literature (see references in the text)

| Signal | Source | OCL | OBL | OBA |
| --- | --- | --- | --- | --- |
| HMGB-1 | aOCY | Recruitment<br>Chemotaxis | Recruitment<br>Chemotaxis | Chemotaxis |
| RANK, OPG<br>M-CSF | OBA<br>aOCY | Recruitment<br>Activation<br>Survival | - | - |
| Sclerostin | OCY | No recruitment | No proliferation<br>No differentiation<br>Apoptosis | No proliferation<br>No differentiation<br>Apoptosis |
| TGF-β<br>BMPs<br>PDGF<br>IGF<br>FGF | OCY<br>OBL<br>OBA | Deactivation | Chemotaxis<br>Recruitment<br>Differentiation<br>Proliferation<br>Survival | Chemotaxis<br>Differentiation<br>Proliferation<br>Survival |

### Signal integration by individual cells. Inhibitory proteins

We now formulate our basic modeling assumptions, which can be summarized as follows. Cells within a BMU integrate signals in their immediate surroundings and the resultant determines a very limited set of actions, namely differentiation, cell division, migration and apoptosis. In addition, we propose that inhibitory proteins, blocking the progression of these actions within each cell type, mediate cell decisions in any bone-remodeling event.

To clarify this last assumption, it is worth recalling the well-known behavior of two inhibitory proteins, Rb and Bcl-2. The retinoblastoma protein (Rb) arrests progression into the cell cycle, whereas the B-cell lymphome-2 protein (Bcl-2) precludes the initiation of the apoptosis program in most cell types, including osteocytes and other stromal cells [36]. More precisely, Rb binds to transcription factors of the E2F family, preventing the progression of the cell cycle to the synthesis stage [40]. When the amount of active Rb falls below a critical threshold, a no-return point is reached (the Restriction Point of the cell cycle) that irreversibly leads to cell division. Analogously, Bcl-2 precludes the release of cytochrome c through the mitochondrial outer membrane, thus avoiding the initiation of the apoptosis program. If Bcl-2 is depleted beneath a suitable value, its inhibitory action is lost and the cell is committed to die [26, 41]. Hence a competition between inhibitory molecules results in a mechanism of cell fate control: the first of these inhibitors (Rb or Bcl-2) that falls below its corresponding threshold value determines the cell decision to divide or die and, importantly, the timing of such decision (see Figure 2.A). This mechanism also provides an explicit link between external signals and cell decisions, since membrane receptors are known to modulate the evolution of inhibitory molecules within the cell [41]. As a matter of fact, it has been recently shown that the interplay between receptor/signal interaction and the internal dynamics of Rb and Bcl-2 suffices to explain the onset of emergent population properties (as clonal expansion and contraction) in the case of immune response to acute infections [42], see also Figure 2.

**Fig 2.**
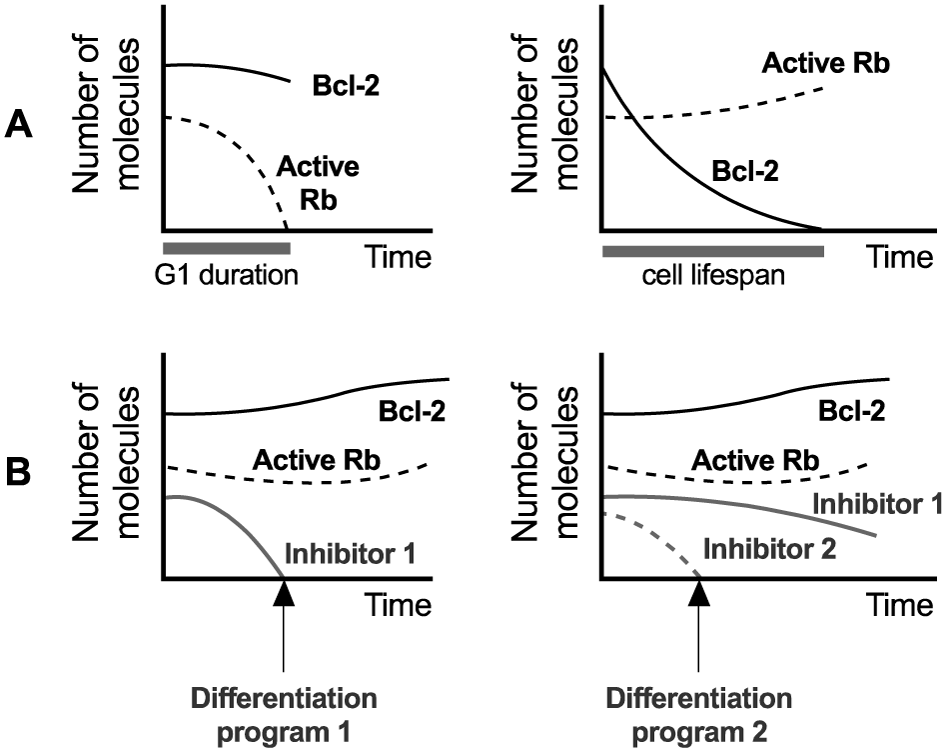
A mechanism of cell fate determination. **A)** Inhibitory proteins Rb and Bcl-2 provide a fate decision mechanism in several cell types. The first of these molecules to reach its critical threshold determines if the cell will divide or undergo apoptosis. For convenience all inhibitory thresholds are set equal to zero. **B)** The presence of inhibitors blocking alternative differentiation programs allows to increase the complexity of this cell fate-decision mechanism. **B Left)** If differentiation is assumed to be blocked by Inhibitor 1, and this inhibitor vanishes before Rb and Bcl-2 do, then the cell will not divide or die, but will instead undergo the differentiation program 1. **B Right)** Two alternative differentiation programs can be controlled by two different inhibitors (1 and 2). If one of them (inhibitor 2 in this case) disappears faster than the remaining molecules, the corresponding differentiation program is selected.

In our case, the occurrence of inhibitory proteins controlling cellular processes during bone remodeling is well documented in the literature. For instance, the roles of Rb, Bcl-2 and the transcription factor Runx2 have been described in [43–45] respectively. Moreover, different restriction points are known to occur for the various cell commitment alternatives involved in bone remodeling. In particular, the onset of two restriction points in the differentiation program of osteoblasts (marking respectively the transition to activated osteoblast and osteocyte types) has been pointed out in [26].

The increased complexity derived from the presence of more than one cell type, together with the existence of several cell fates (division, apoptosis or alternative differentiation programs) introduces new possibilities with respect to the basic dichotomy cell division vs. cell death considered in [42]. However, the underlying logic can be extended to account for these new alternatives. For instance, several inhibitory molecules can simultaneously block the progression of alternative differentiation programs. In this case, the first inhibitor to reach its critical threshold will determine the fate of the cell (see Figure 2.B). We propose that cell choices thus determined are mutually exclusive. This seems to be the case for BMU cells. For instance, osteoblasts that start the differentiation program or decide to secrete osteoid matrix do not complete the cell cycle, and therefore do not proliferate [26]. We also remark that this mechanism allows for one signal to trigger alternative cell decisions depending on its concentration.

Bearing in mind the multiplicity of signals recalled in Table 1, as well as the redundancy often observed in their functioning, we propose that the effective operation of a BMU can be modeled by means of a reduced version of the complex signaling network previously sketched. Specifically, we propose that three cell-released signals, denoted by R, S and T (with analogue roles to those of RANK, sclerostin and TBF-*β* respectively; see Table 1) plus one osteoclast cue (see [34]), that keeps such cells moving towards intact bone, acting on three types of internal inhibitors suffice for that purpose. The effect induced by such signals in the cell types involved is described in Table 2.

**Table 2.** Generic signals to be considered in our model and a list of the actions they induce on cell types of a BMU.

| Signal | Source | OCL | OBL | OBA |
| --- | --- | --- | --- | --- |
| R | OBA | Activation<br>Survival | - | - |
| S | OCY | No recruitment | No proliferation<br>No differentiation<br>Apoptosis | No proliferation<br>No differentiation<br>Apoptosis |
| T | OCY<br>OBL<br>OBA | Deactivation | Proliferation<br>Differentiation<br>Survival | No proliferation<br>No differentiation<br>Apoptosis |

### A model for BMU operation

We next describe the cell algorithms that constitute our proposed model. For simplicity, we will consider a two-dimensional cross section of bone adjacent to a section of bone marrow. The bone section considered will be thought of as a lattice with coordinates x, y, divided in boxes of equal size. Any such box can either remain empty or be occupied by only a single cell. On such region we will define a cellular automaton (CA) to describe the dynamics of the remodeling process. To that end, we will implement cellular algorithms based on the biological assumptions stated below. A first assumption is that progression within the cell cycle, apoptosis and the initiation of differentiation programs are initially blocked by specific inhibitory molecules as described above. This would allow for newly formed cells to remain in the first stage of the cell cycle (G1) before choosing a given cell fate. During this stage, membrane receptors interacting with external signals govern the dynamics of inhibitors, thus controlling the eventual decision of the cell. The effect of the complexes formed by membrane receptors and external cues in the inhibition (resp. activation) of any cell fate choice takes place through an increase (resp. decrease) in the amount of the inhibitory molecule controlling this choice. Cell fate is decided when the concentration of the first of the inhibitors considered attains its threshold value, that we set equal to zero for simplicity.

In our case, the inhibitor dynamics will be assumed to be of the following form:

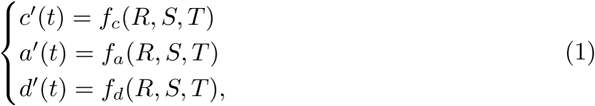

where *c*(*t*), *a*(*t*) and *d*(*t*) respectively denote the concentrations of division, apoptosis and differentiation inhibitors at time *t* and *f_c_*, *f_a_*, and *f_d_* are functions of three external signals *S*, *R* and *T* (see figure 3). For simplicity, we shall assume in the sequel such functions to be piecewise linear. We next describe the main details of the decision algorithm proposed to describe BMU operation.

**Fig 3.**
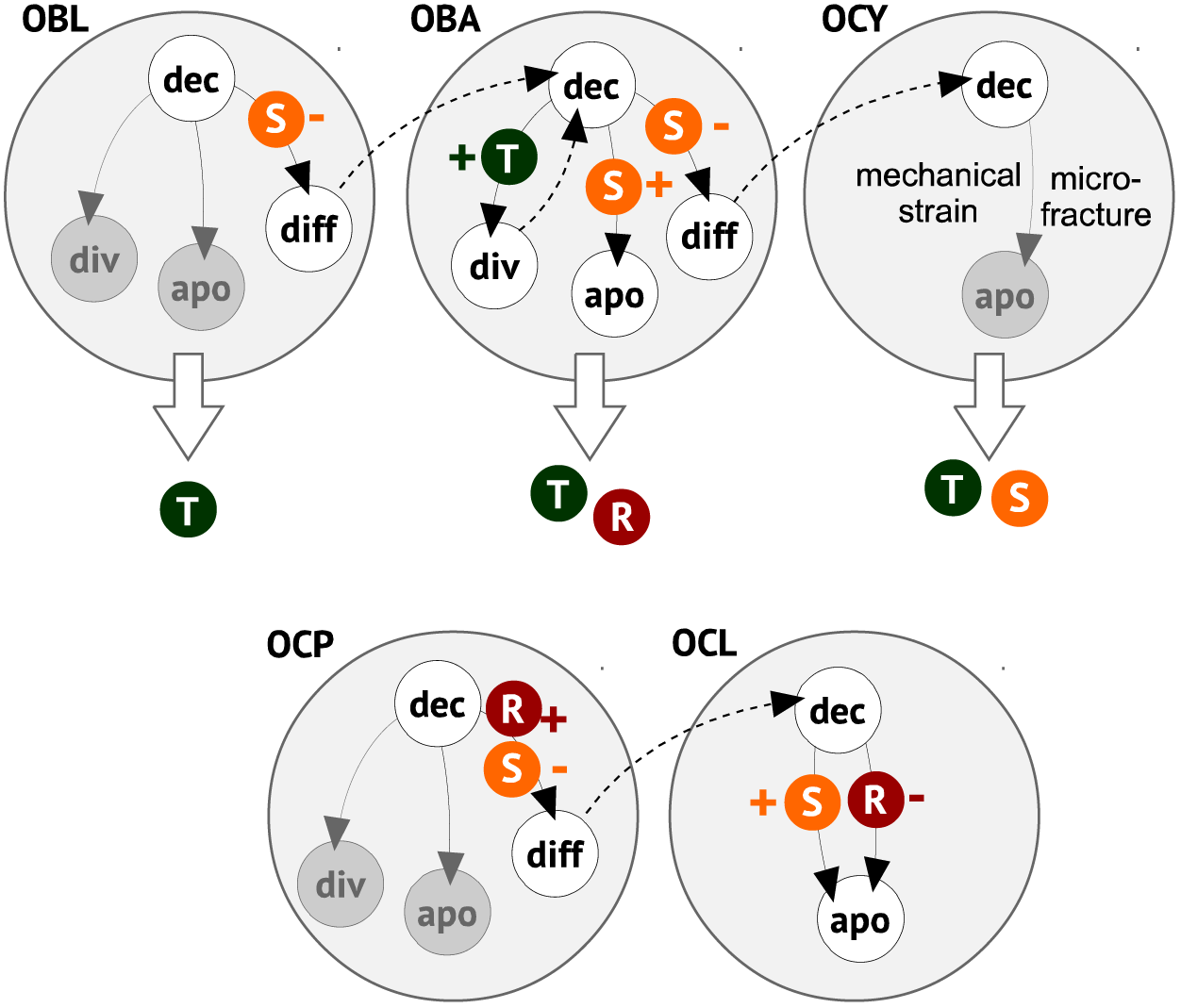
A minimal model of BMU software. Signals *R*, *S* and *T* (in small circles) are produced by the cell types listed. They may inhibit (−) or induce activation (+) on the actions considered. Activation results from double inhibition, that is by lowering the concentration of an inhibitor. Within any cell type, possible decision choices are indicated by thin arrows. Dashed arrows correspond to a starting decision stage in a newly formed cell. Extracellular release of signals by any cell type (for instance, *R* and *T* in the case of active osteoblasts) is denoted by thick, white arrows.

### Osteoblasts (OBL)

Osteoblasts (OBL) are initially located at the interface between bone matrix and bone marrow. In normal conditions, osteoblasts homeostasis is maintained by a continuous cell turnover, involving both division and apoptosis [46]. When a remodeling process starts, OBLs can choose among three alternative programs: division, apoptosis and differentiation into active osteoblasts (OBAs). We shall assume that OBL division an apoptosis just balance each other, and therefore focus on the third choice above. Activation is known to be mainly blocked by the release of sclerostin by osteocytes [6, 10]. This will be modeled by assuming that signal *S*, produced by osteocytes, increases the amount of a differentiation inhibitor in OBLs, denoted by *d_B_*, according to the following dynamics:

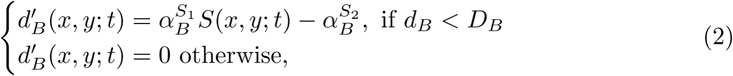

where prime denotes time derivative, (*x*, *y*) denotes the position of the OBL, *D_B_* is the maximum amount of inhibitor that can accumulate within a OBL *S*(*x*, *y*; *t*) is the amount of signal *S* at location (*x*, *y*) at time *t* and 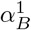 and 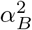 are positive structural parameters. During a bone remodeling process, osteoblasts remain in their original positions, delimiting the BRC [11]. Accordingly, OBLs do not move in our model. We recall that differentiation of a particular OBL is blocked while *d_B_ >* 0, and that the process of differentiation is triggered by the condition *d_B_* = 0.

### Active osteoblasts (OBA)

Once differentiated, OBLs become active osteoblasts (OBA). OBAs have three possible choices: cell division, cell death and differentiation into osteocytes (OCY). Cell division proceeds according to the following rules:

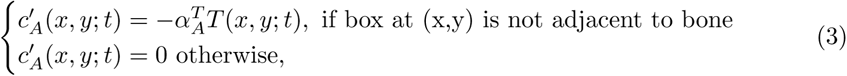

where (*x*, *y*) denotes the position of the OBA, *T* (*x*, *y*; *t*) is the amount of signal *T* at location (*x*, *y*) at time *t* and 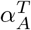 is a positive parameter. Cell division only occurs when the condition *c*(*A*) = 0 is verified. Notice that, in our model, the movement of active osteoblasts front is just a consequence of cell division.

When OBAs are attached to the bone surface, cell division is no longer possible for lack of space. In that case OBAs secrete osteoid matrix; this last is represented by an order parameter *b*(*x*, *y*; *t*) which ranges from 0, corresponding to eroded bone, to value 1 for fully functional bone. Apoptosis, differentiation into OCY and osteoid production can take place, according to these equations:

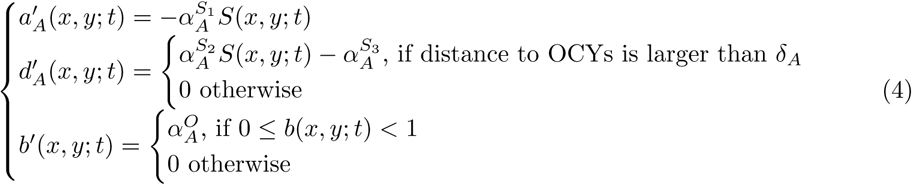

where *S*(*x*, *y*; *t*) is the amount of signal *S* at location (*x*, *y*) at time *t*, 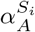 (for *i* = 1, 2, 3) are positive structural parameters, 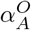 the rate of bone production, and *δ_A_* is a positive constant that represents a threshold in cell-to-cell contact inhibition by OCYs on OBAs differentiation, resulting from gap junction communications between cells [10, 13, 47].

### Osteoclast precursors (OCP)

OBA can recruit OCP to the bone remodeling zone by secreting signal *R*, while signal *S* inhibits OCP recruitment. For simplicity, we will not consider in the model OCP division and apoptosis, and assume instead that enough OCP are available in the bone marrow (in particular, in the locations adjacent to those occupied by OBAs). OCPs attach to the bone surface upon their recruitment by OBAs, after which they become active osteoclasts (OCL). We assume that OCP activation is blocked by inhibitor *d_P_*, whose dynamics is modeled as follows:

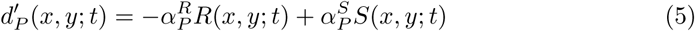

where *R*(*x*, *y*; *t*) and *S*(*x*, *y*; *t*) are respectively the amounts of signal *R* and *S* at location (*x*, *y*) at time *t* and 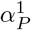 and 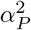 are positive parameters.

### Osteoclasts (OCL)

Once recruited to the BRC, OCPs become active osteoclasts (OCL) and start eroding the bone matrix, thus leading to the formation of a cutting cone [34]. OCLs are assumed to be endowed with one apoptosis pathway controlled by signal *S*, produced by OCYs. This signal is instrumental in determining the depth that will be reached by the cutting cone. In particular, OCLs in locations with high concentrations of *S* will stop digging into the bone matrix and die. We will model the apoptosis pathway controlled by signal *S* by assuming that cell death is blocked by inhibitor *a_C_*, whose dynamics is given by the

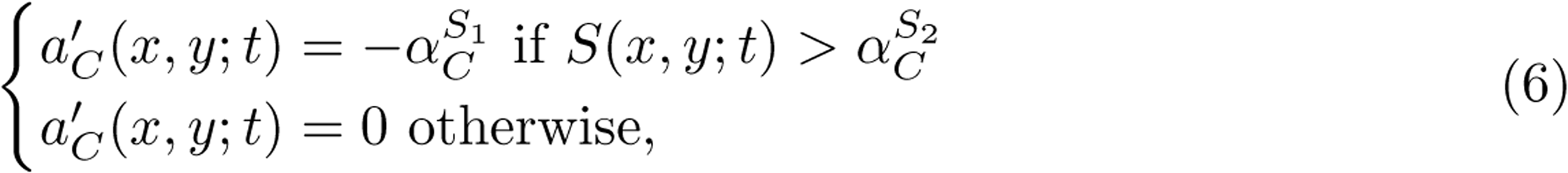

where *S*(*x*, *y*; *t*) is the amount of signal *S* at location (*x*, *y*) at time *t* and 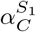 and 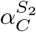 are positive parameters.

We next describe the movement of osteoclasts during a bone remodeling event. We will assume that osteoclasts remove bone at a rate that depends on the amount of signal *S* and move to an adjacent box, along the normal to the ossification front, once they have removed bone in their current position. Bone resorption will be modeled by means of the following equation:

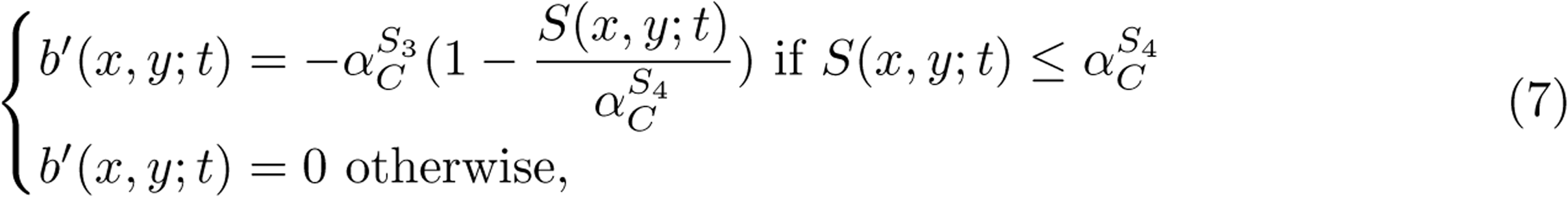

where *b* measures the presence of bone, so that *b* = 1 corresponds to functional bone, and *b* = 0 to disposed bone. *S*(*x*, *y*; *t*) is the amount of signal *S* at location (*x*, *y*) at time *t* and 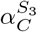 and 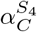 are positive parameters.

### Signal diffusion and decay

The dynamics of signals in the CA is now described as follows. At any given time, they are produced by the corresponding cell types at constant rates. Signals are also assumed to undergo Arrhenius-type decay (meaning that their decay rate is proportional to their concentration). Signal transport through the bone matrix is conveniently represented as a diffusion process. We assume that such diffusion occurs faster that internal cell processes. In order to account for these two time scales simultaneously, we model diffusion by calculating, at each box and each iteration step of the model, a weighted average of the amount of signals in neighboring positions (see Figure 4.A). Cells decisions are made upon comparing the amount of different signals in their adjacent boxes.

**Fig 4.**
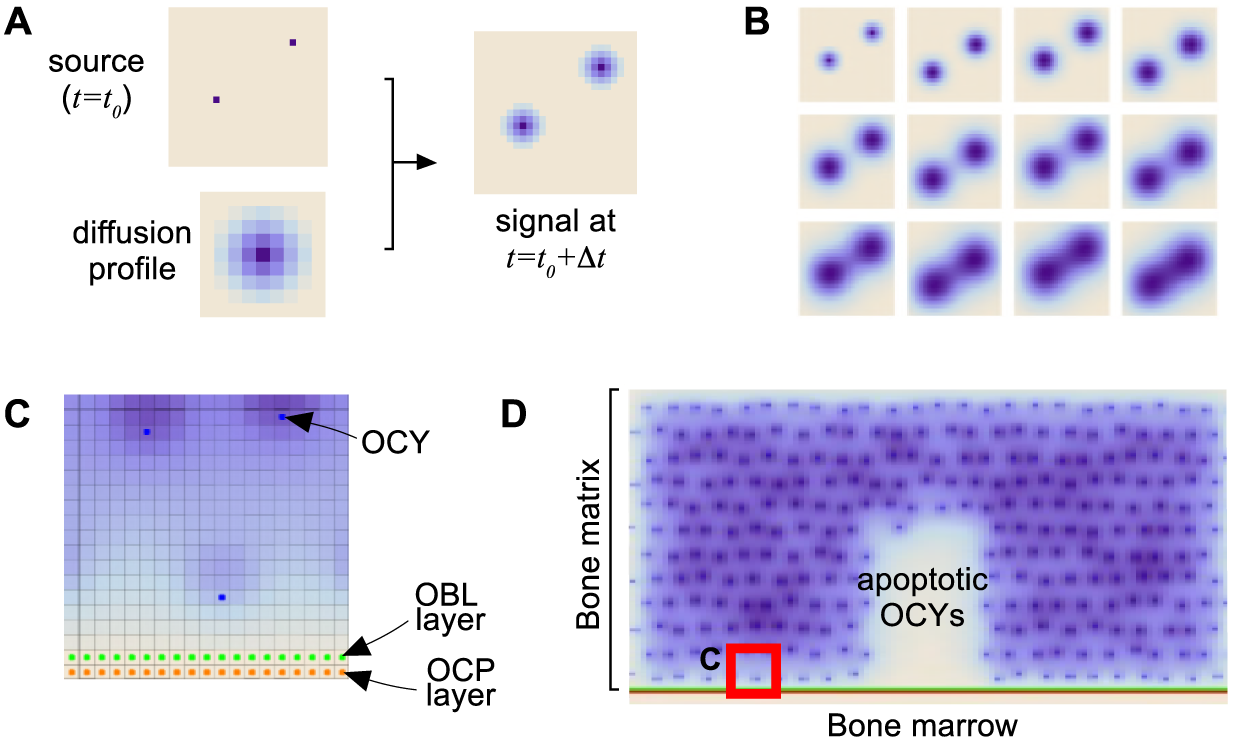
Spatial aspects of the model. **A)** Signal diffusion takes place at a different scale than cell decisions. Let ∆*t* be the characteristic time step for cell processes. In order to model how signals spread out, we calculate the diffusion profile created by the signal release from each cell during this time interval. Diffusion can then be modeled at this characteristic time by applying the corresponding profile to any source of signals in the CA. **B)** Snapshots of a signal diffusion from the source shown in A. Time increases from left to right and from top to bottom. **C)** Spatial arrangement of the main elements of the model. Bone matrix is locally represented by means of a lattice consisting of equal-size boxes, each able to accommodate one cell. The lower boundary of the box considered in C) is represented as lined with an osteoblast layer, lying upon a layer of osteoclast precursors. **D)** Initial configuration of the model. At a selected place within the bone matrix, osteocytes (blue dots) may undergo apoptosis, thus triggering BMU operation in the corresponding region (blank). The lattice element considered in C) is represented here for comparison purposes (red square in the lower left corner). Shades of blue represent different concentrations of signals in each box.

### BMU initialization

As a starting point we consider a population of osteocytes (OCY), regularly distributed within the bone matrix, and a layer of osteoblasts lining the bone section (see Figure 4.C). Simulations of the model are started after apoptosis of a group of osteocytes has occurred (see Figure 4.D).

## Results

In this Section we present some results obtained upon simulation of the model whose elements have been described in our previous Section. To keep the flow of the main arguments here, we postpone some technical details to the Supporting Information at the end of the paper. Specifically, the reader will find there a flow chart describing the order in which the corresponding algorithms are applied, as well as a list of the parameters used in the simulations whose results will be shown below.

## The BMU software proposed successfully recapitulates bone remodeling

Upon induction of osteocyte death in a region, the proposed cell algorithms provide a mechanism of bone remodeling which sequentially reproduces the main phases of the actual process. See Figure 5 and its caption for more details. A video corresponding to this simulation is provided with the Supporting Information.

**Fig 5.**
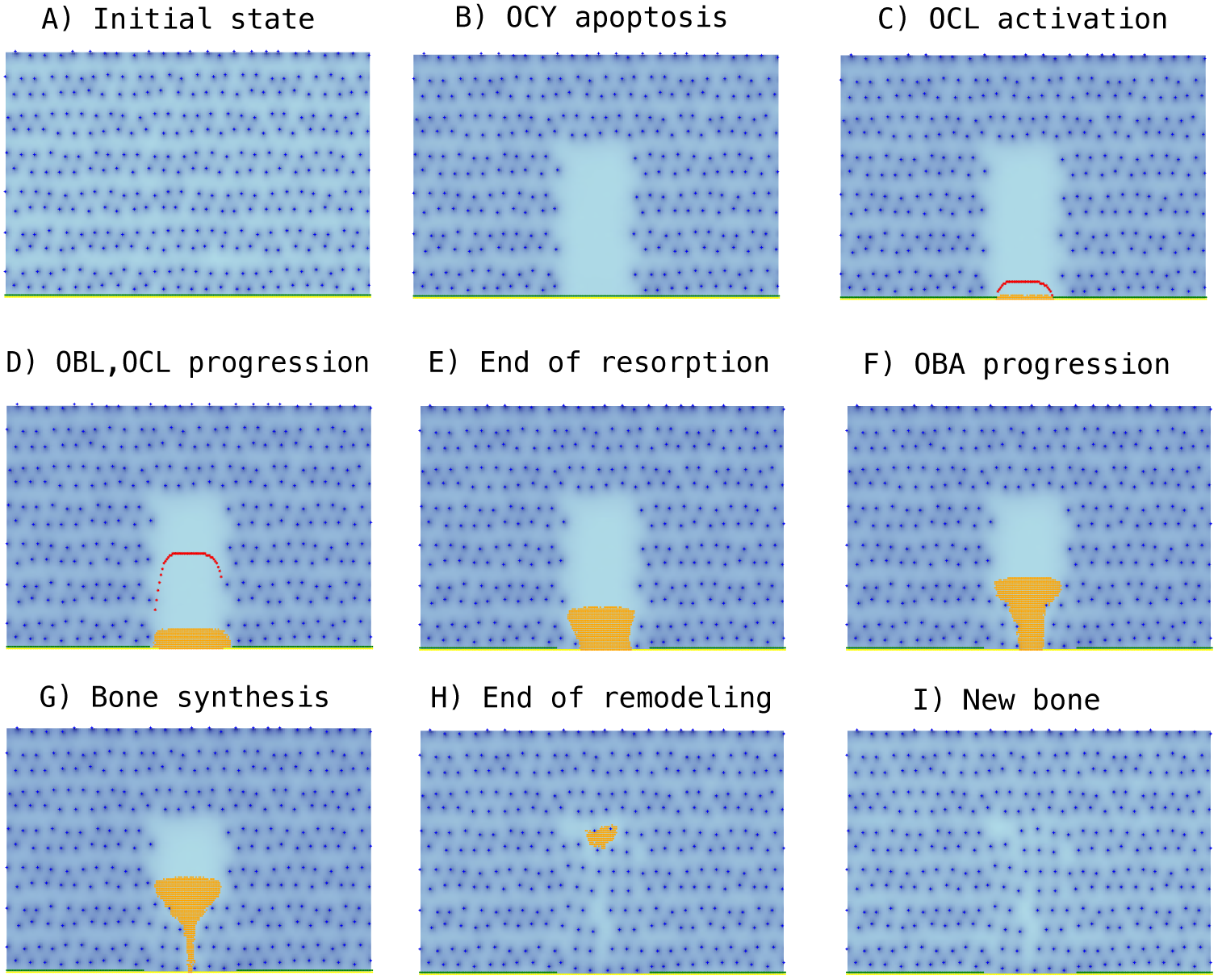
Snapshots corresponding to sequential stages in a bone remodeling event. **A**) A planar bone region is considered where active osteocytes (OCY, blue dots) are regularly distributed. Osteoblasts (OBL) are located at the lower side, which represents the interface between bone and bone marrow. **B)** OCY apoptosis is induced in a subset (in light blue) of the previous region, which leads to a drop in OCY inhibitory action in that place. **C)** Lack of enough osteocyte inhibition results in OBL activation, and the recruitment of osteoclast precursors which differentiate into active osteoclasts, OCL (in red). **D)** OCL form cutting cones that move into the apoptotic region destroying old bone (bone resorption). **E)** OCL-driven resorption proceeds until the inhibitory effect of the remaining osteocytes precludes further OCL progression. **F)** Activated osteoblasts (OBA, in orange) move in the wake of OCL. **G)** Upon successive rounds of cell division, an increasingly larger region of OBA is filling the region previously occupied by decaying bone, expanding behind the OCL-lined cutting cone boundary where they secrete osteoid matrix. Part of them subsequently differentiate into OCY (blue dots). **H)** When remodeling is finished, a new distribution of osteocytes is achieved in the remodeled region, and **I)** new bone is eventually formed.

We point out that simulation of our model reveals that both the initial state of fully homeostatic bone, and the final stage after bone remodeling, remain stationary in time. In other words, running the model on such configurations does not result in any noticeable change in such states.

## BMU algorithm is robust to variations in the size of the BRC region

The model proposed can be run when OCY apoptosis is assumed to occur in regions of different sizes. In each case, the area of the corresponding Bone Remodeling Compartment (BRC) is shown to depend on the extent of the induced OCY apoptosis. See Figure 6.

**Fig 6.**
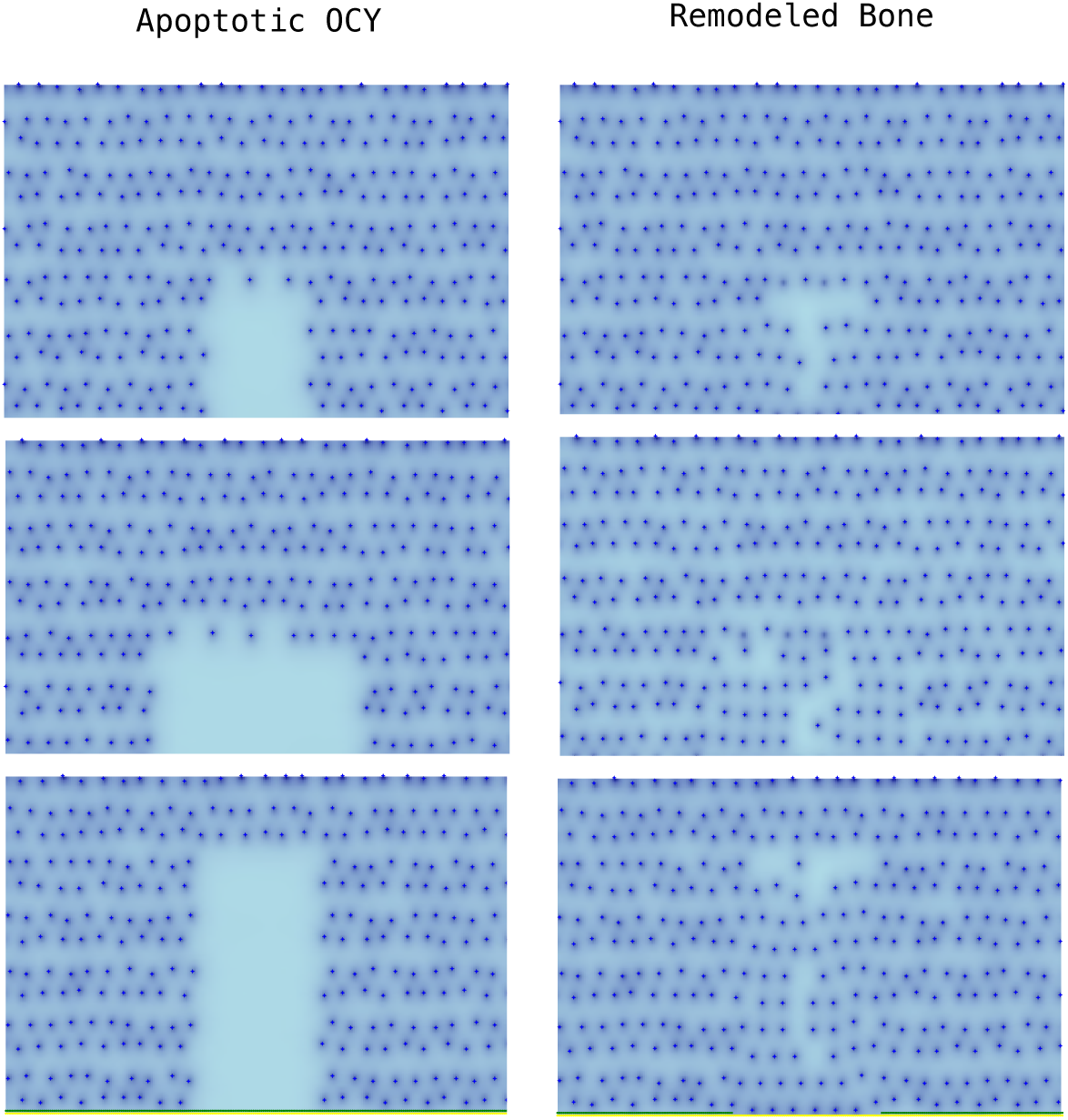
Simulation of three different scenarios of OCY apoptosis. Images correspond to results obtained in each case after the simulation of the model since OCY apoptosis until the end of the remodeling process. The shape and size of the resulting BRC is determined in each case by the region where OCY apoptosis has occurred.

## Osteoclast lifespan depends on the depth of the BRC

A prediction of the model is that average osteoclast lifespan is not a priori determined, but depends instead on the size of the region where bone remodeling has to be performed. This fact is illustrated in Figure 7 below, where mean OCL lifespan is plotted in terms of BRC depth.

**Fig 7.**
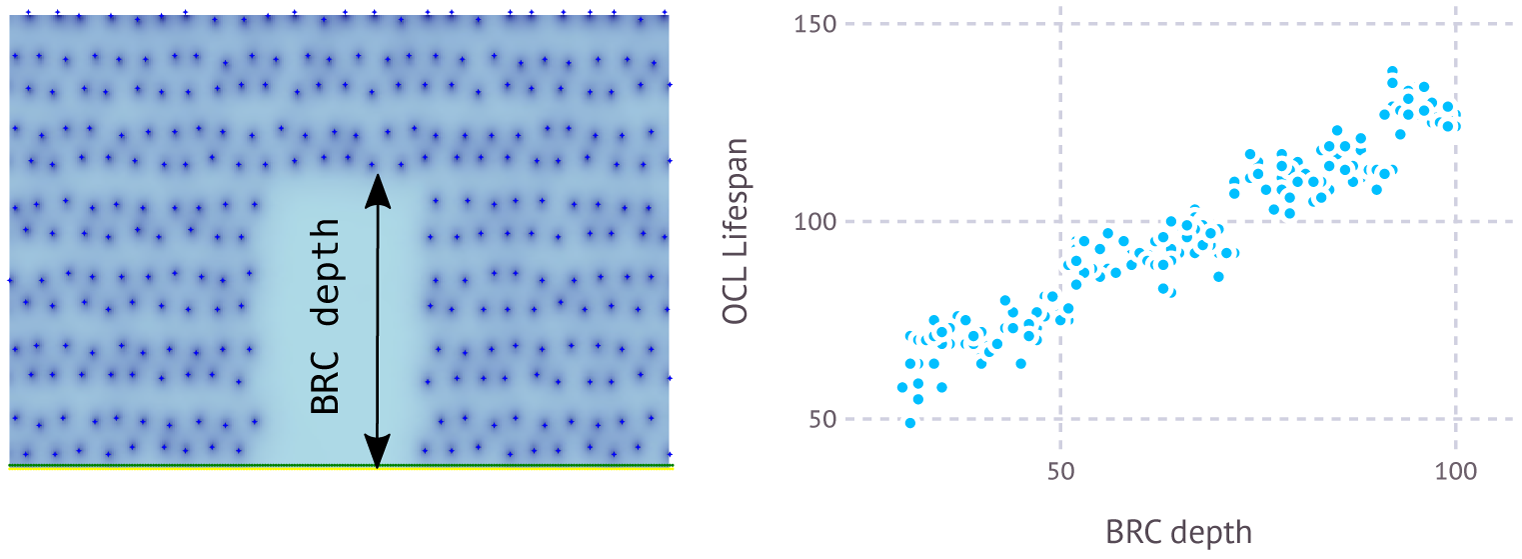
Dependence of OCL lifespan on BRC size. **Left)** A region where bone resorption has occurred following OCY apoptosis is depicted. BCR depth is defined there as the longest distance within that region measured perpendicular to its lower side. **Right)** A plot of OCL lifespan during bone remodeling events in terms of BRC depth obtained after simulations of the model (each dot corresponds to a simulation). Note that OCL live longer when BRC is larger.

## The BMU software is robust to signal noise

As we have seen above, cell dynamics within a BMU depends on cell-to-cell communication mediated by signals that diffuse through the bone matrix. The coupling of old bone resorption and new bone formation can therefore be affected by signal noise due, for instance, to bone matrix spatial heterogeneity. In Figure 8 we show that the suggested BMU software is robust to environmental noise.

**Fig 8.**
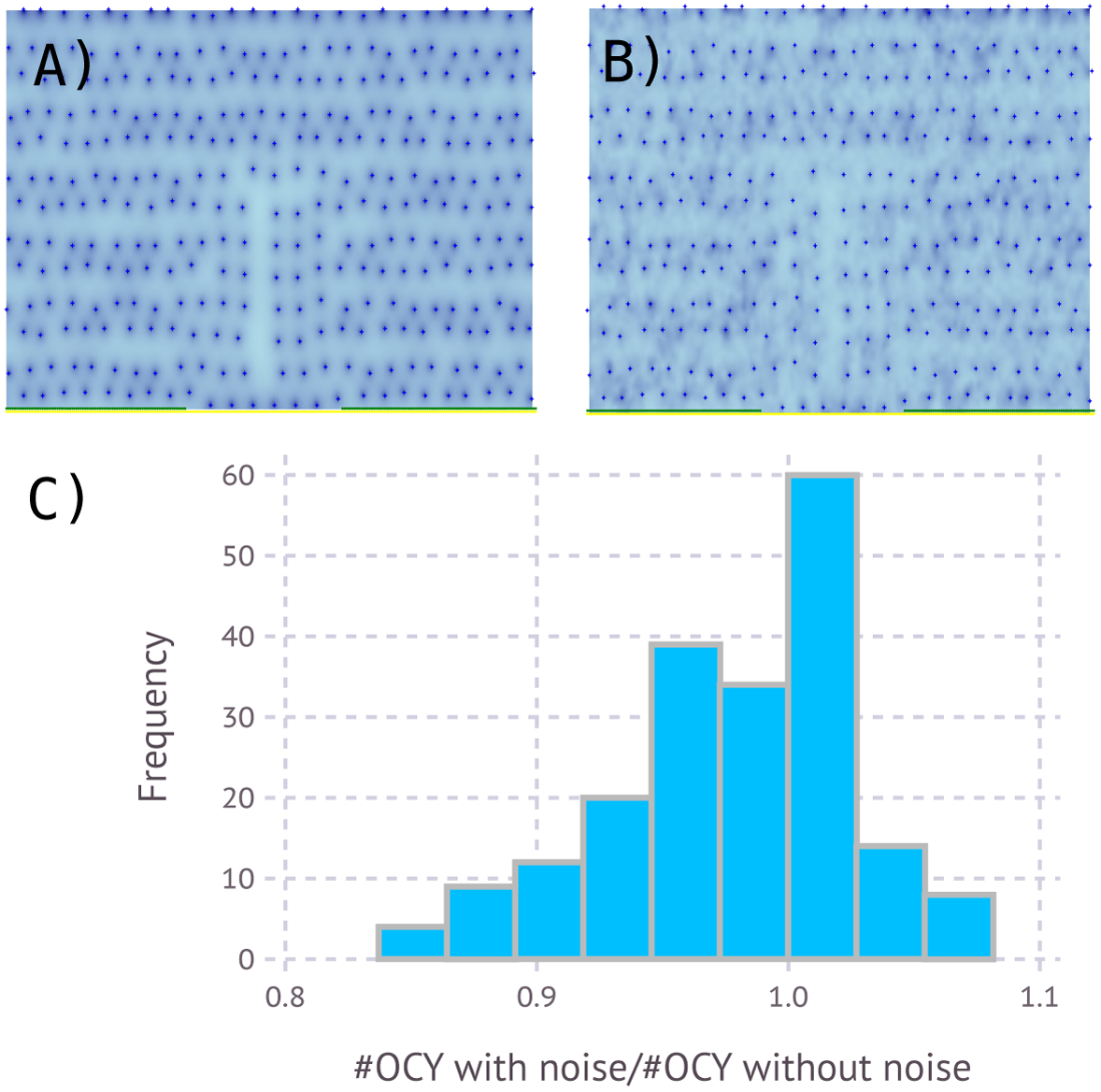
Effect of signal noise in BMU operation. **A)** Simulation corresponding to a bone remodeling without noise in signal S diffusion. The picture corresponds to the new bone after remodeling. **B)** In spite of noise in signal S, the new distribution of OCY is similar to A. **C)** Histogram of the ratio between the number of newly formed OCY in the BRC after bone remodeling with and without noise (the result was obtained after 200 simulations of the model).

## Discussion

In this work we have proposed and analysed a minimal model for bone remodeling carried out by Bone Multicellular Units (BMUs) to replace old bone by new one, a process which is often triggered by changes in mechanical load. We have already recalled that such remodeling is continuously taking place during human life, so that on average the whole skeleton is renewed approximately every 10 years [14]. BMUs activity is usually carried out on small regions, of about 4000 microns in width [9], and therefore the total number of cells involved in each remodeling event is comparatively small. The previous length scale can also be used to characterize micro-fractures (arising for instance from physical exercise) which are also self-repaired in the same manner. BMUs operation can thus be viewed as the basic building block to keep homeostasis in bone tissue. When normal function in bone is compromised by large-scale hazards (say, fractures), more sophisticated repair mechanisms, involving in particular inflammatory signals, blood clot formation and the production of different callus templates for bone repair are involved. These mechanisms have not been discussed here.

Our goal in this work is to identify, by means of a mathematical model, a simple algorithm for BMU operation. To do this, we have kept to a few basic principles. First, at any time during that process, each cell within the different lineages present in a BMU should individually select one among a restricted set of actions: divide, die, migrate or differentiate. Such choice is not assumed to be random, but determined by feedback from the surrounding medium into the internal dynamics of molecules which act as decision inhibitors. No previous planning is presupposed, genetically or otherwise. Second, only a minimal number of signals are selected that allow for BMU functioning.

As recalled in a previous Section, molecular cues acting as those retained in our model have been already described. In particular, cell decision inhibitors have been identified, and effects induced by the signals retained have been shown to be caused by chemical mediators experimentally detected. However, we have not aimed at producing a comprehensive model where all data currently known could be fitted. We have instead attempted to show what the basic ingredients of an operational BMU could be like. Implicit in this approach is the assumption that some signals already identified are multi-functional (thus yielding different effects in different cell lines) and that in general some signaling networks observed can be redundant, perhaps as a consequence of having been arrived at through different paths in the course of evolution. Notice, however that we do not claim that our proposal is the only possible one. In principle, there could be alternative minimal models for what we have called a BMU software. We believe however, that the degree of complexity retained in our model could not be significantly reduced by alternative mechanisms.

Selecting a minimal model has the advantage of dealing with a comparatively small number of parameters. Concerning this last issue, we wish to emphasize that no parameter-fitting attempt has been made here. Our concern is to show that the model proposed has not only the potential to reproduce standard features in bone remodeling, but also to draw conclusions about the manner in which this process occurs that could not be a priori anticipated before simulating it. Accordingly, we will be content to select a set of coefficients for which such goals are achieved. Parameters appearing in the actual BMU software should be of a structural nature, and their precise values have to be experimentally determined. In fact, the circuitry underlying BMU operation is expected to be rather sensitive to small variations in such parameter values, a fact repeatedly observed in processes which have been arrived at in the course of evolution. An example of such situation is provided by the Krebs cycle [48, 49] which consists in a network of biochemical reactions taking place exactly at very precise rates and proportions. On the other hand, it is well known that a given phenomenon can be fitted by utterly different models, relying on different principles, upon appropriate choices of a suitable (and often rather modest) number of parameters appearing in them [50, 51]. We have shown herein that the parameters retained in our model can be selected so that BMU standard operation is reproduced and, importantly, consequences such as the dependence of osteoclasts lifespan on the size of the region to be repaired can be inferred that were not a priori clear before the model was simulated. We do not claim, however, that the results obtained provide a validation of the model. In our opinion, such validation could only stem from experimental determination of such values.

We conclude this discussion by briefly remarking on possible future directions arising from this work. To begin with, the experimental question of determining the operational parameters introduced in our model should be again stressed. If and when this is done, we would have in place a BMU chip structure to be used as a building block for operational purposes. On the other hand, one could simulate on models as that presented here the impact of external cues in the modulation of the repair process, either to slow it down or to accelerate it. Finally, as the size of the region to be repaired increases, a threshold should be reached beyond which inflammatory signals become relevant, and a new and more complex stage is entered. Understanding the matching between these two cases could be instrumental in designing techniques to foster bone fracture repair. We intend to pursue some of these subjects elsewhere.

## Supporting Information. Details of computer simulations of the model of BMU operation during bone remodeling

### Time evolution of the BMU

#### Initialization

Simulations take place in a two-dimensional cellular automata (CA) of 150 × 250 boxes. 125 rows of this matrix are initially occupied by bone matrix and the rest by bone marrow. A row of OBL lies in the interface between them. OCYs are initially located at fixed, regularly spaced positions within the bone matrix. A random perturbation is then applied to their positions to avoid any potential effect of symmetries on the simulations of the model. At this point, OCYs and OBLs release signals that diffuse through the bone matrix until an equilibrium is attained. This equilibrium is determined by the balance between signal production and decay. Periodic boundary conditions are considered on both lateral sides of the region considered, and zero signal flux is imposed on the top and the bottom. A region of OCY apoptosis is randomly defined within *a priori* defined boundaries (*Depth*:= maximum depth and *Width*:= maximum width).

#### Time evolution

Simulations of the model occur in discrete time steps of length Δ*t* = 0.01. The input for an iteration of the model corresponding to time *t* = *t*_0_ includes a set of vectors (one for each of the cells coexisting in the model at that time) with the coordinates of the box occupied by the cell, and its state variables (amount of inhibitory molecules).

##### 1. OBL

The state variable considered in the model for the *i*-th OBL is the amount of activation inhibitor *d_Bi_*. The corresponding vector is:

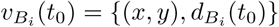

##### 2. OBA

For each OBA there are three different inhibitory molecules, namely *c_A_*, *d_A_* and *a_A_*, blocking cell division, differentiation and apoptosis. The state vector for the *i*-th OBA is:

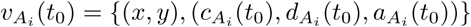

##### 3. OCP

Differentiation of OCP into OCL is controlled by the inhibitor *d_P_*, so that the state vector of the *i*-th OCP is given by:

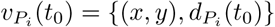

##### 4. OCL

Active osteoclasts dig into old bone while alive before they die by apoptosis. The state vector for the *i*-th OCL includes the amount of apoptosis inhibitor *a_Ci_*:

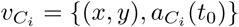

Input also includes the values of bone density *b* and signals *S*, *T* and *R* at each box of the CA. The implementation was carried out with two different tools. The Mathematica platform (Wolfram Research, Inc., Mathematica, Version 9.0, Champaign, IL) was used to obtain Figure 4. The remaining figures and simulations were carried out with the open source *Julia Language* [52], which is endowed with internal mechanisms as *multiple dispatch* and JIT compilation that allowed us to significantly reduce the computational time.

Each iteration starts with the update of cell internal inhibitors according to the equations that describe the dynamics of the state variables (see Models Section). If neither the amount of bone at a box occupied by a OCL or some of the inhibitors goes to zero during a time step, the model moves to the next time step. Alternatively, the model displays different behaviors depending on what variable (or variables) have gone to zero:

##### 1. OBLs activation inhibitor (*d_B_*)

This corresponds to the activation of one or more OBLs. In this case, the activated OBL are removed from the list of cells of the model, and their positions in the CA are occupied by newly formed OBAs.

##### 2. OBAs division inhibitor (*c_A_*)

OBAs form a growing front of cells that progresses by division behind the cutting cone created by OCLs. By construction, only OBAs that are not located in boxes adjacent to bone can divide. Moreover, OBA division can only occur if some of the neighboring positions is free. In this case, the position of a new OBA is selected at random from the free neighbors in the CA.

##### 4. OCPs differentiation inhibitor (*d_P_*)

OCPs are located adjacent to the layer of OBLs. In case of activation, the corresponding OCP is replaced by a newly formed OCL.

##### 5. OCLs apoptosis inhibitor (*a_C_*)

OCLs whose apoptosis inhibitor disappears are removed from the set of cells in the model for the next iteration.

##### 6. Bone removal by OCPs (*b_P_*)

OCLs remove old bone from the box where they are located according to equation (7).When the amount of bone goes to zero, the OCL moves to one of the unoccupied adjacent boxes in the bone matrix. The OCL chooses the closest box that is farther from the OBL layer. OBAs deposit osteoid matrix at a fixed rate in positions attached to bone, and where bone was previously removed from the box. After cell fate choices, the set of cells is updated. Cell produce signals which subsequently diffuse and decay. The model starts a new iteration.

#### End of the simulation

Simulations end when the bone remodeling process is completed (old bone has been replaced by new osteoid matrix) and a new equilibrium (for signals *S* and *T*) has been reached. A video showing one sample of the simulation (from initial OCY apoptosis to final equilibrium and renewed bone) can be downloaded from: https://drive.google.com/file/d/144AT36uRrv1IZMFXLvNmulUmXQVaRbEl/view?usp=sharing. The movie should be played with ADOBE ACROBAT READER®.

~~~
**Pseudo-code guidelines**
   **Input parameters**
       *t_max_*:= maximum duration of simulations
       *Depth*:= maximum depth of the apoptosis region
       *Width*:= maximum width of the apoptosis area
       *bone*(*x*, *y*):= 1 if the box is occupied by bone at initial time and 0 otherwise
       *position*(*OCY,* 0):= initial position of OCYs
       *position*(*OBL,* 0):= initial position of OBLs
       *position*(*OCP,* 0):= initial position of OCPs
   **Structural parameters**
       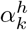:= effect of external signals of type *h* on the dynamics of inhibitors in cells of type *k*
       *D_B_*:= maximum amount of inhibitor *d_B_* that can accumulate inside a OBL
       *d_ph_*(*d*):= diffusion profile of signal *h* as a function of distance *d*
       *γ_h_*:= rate of decay of signal of type *h*
   **Variable parameters**
       *{a_k_*_0_*, d_k_*_0_*, c_k_*_0_*}*:= amount of inhibitors in newly formed cells of type *k δ_A_* Distance of OCY inhibition of OBA differentiation
       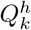:= rate of production of signal of type *h* by cells of type *k*
   **Initialization**
     1 Input the bone matrix *bone*(*x*, *y*)
     3 Input the the locations of OCY, OBL and OCL.
     4 Production, diffusion and decay of signals *S* and *T* until equilibrium
     5 Define a region of OCY apoptosis
     6 Set *t*:= 0
     7 **REPEAT**
         8 Input the set of positions and state variables of all cells
         9 Input the amount of signals *S*, *T* and *R* at each box of the CA
         10 Integration of the equations for the internal inhibitors for a time step Δ*t*
         11 If (*d_Bi_*(*x*, *y*; *t*) 0) then remove the *i*-th OBL and add a new OBA at box (*x*, *y*)
         12 If (*c_Ai_*(*x*, *y*; *t*) *≤* 0) then
             13 add a new OBA at a box adjacent to (*x*, *y*)
             14 recalculate OBAs positions
         15 If (*a_Ai_*(*x*, *y*; *t*) *≤* 0) then
             16 remove the *i*-th OBA
             17 recalculate OBAs positions
         16 If (*d_Ai_*(*x*, *y*; *t*) *≤* 0) then
             17 remove the *i*-th OBA
             18 add a new OCY at (*x*, *y*)
             19 recalculate OBAs positions
         20 If *d_Pi_*(*x*, *y*; *t*) *≤* 0 then
             21 remove the *i*-th OCP
             22 add a new OCL at (*x*, *y*)
         23 If *a_Ci_*(*x*, *y*; *t*) *≤* 0 then remove the *i*-th OCL
         24 If *b_Bi_*(*x*, *y*; *t*) *≤* 0 then
             25 set *bone*(*x*, *y*) = 0
             26 move the OCL to a new position in the bone matrix
         27 Cell production, diffusion and decay of signals *S*, *T* and *R* during the time step Δ*t*
         13 set *t* = *t* + Δ*t*
             # We remark that several cell decisions can take place simultaneously
     32 **UNTIL** the end of bone remodeling process
~~~

**Fig A.3.**
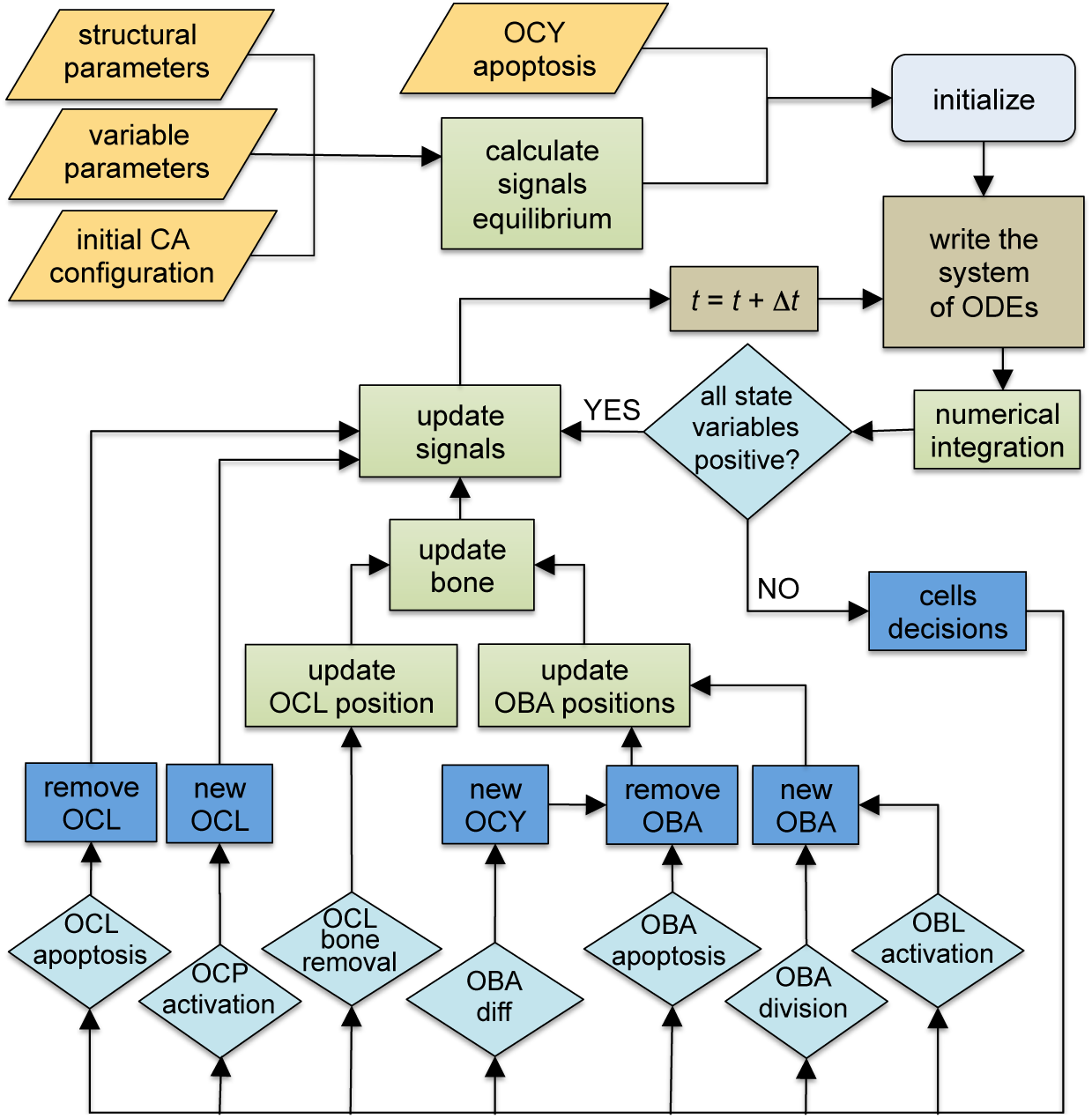
Flowchart of the proposed model of BMU software.

**Table A.1.**
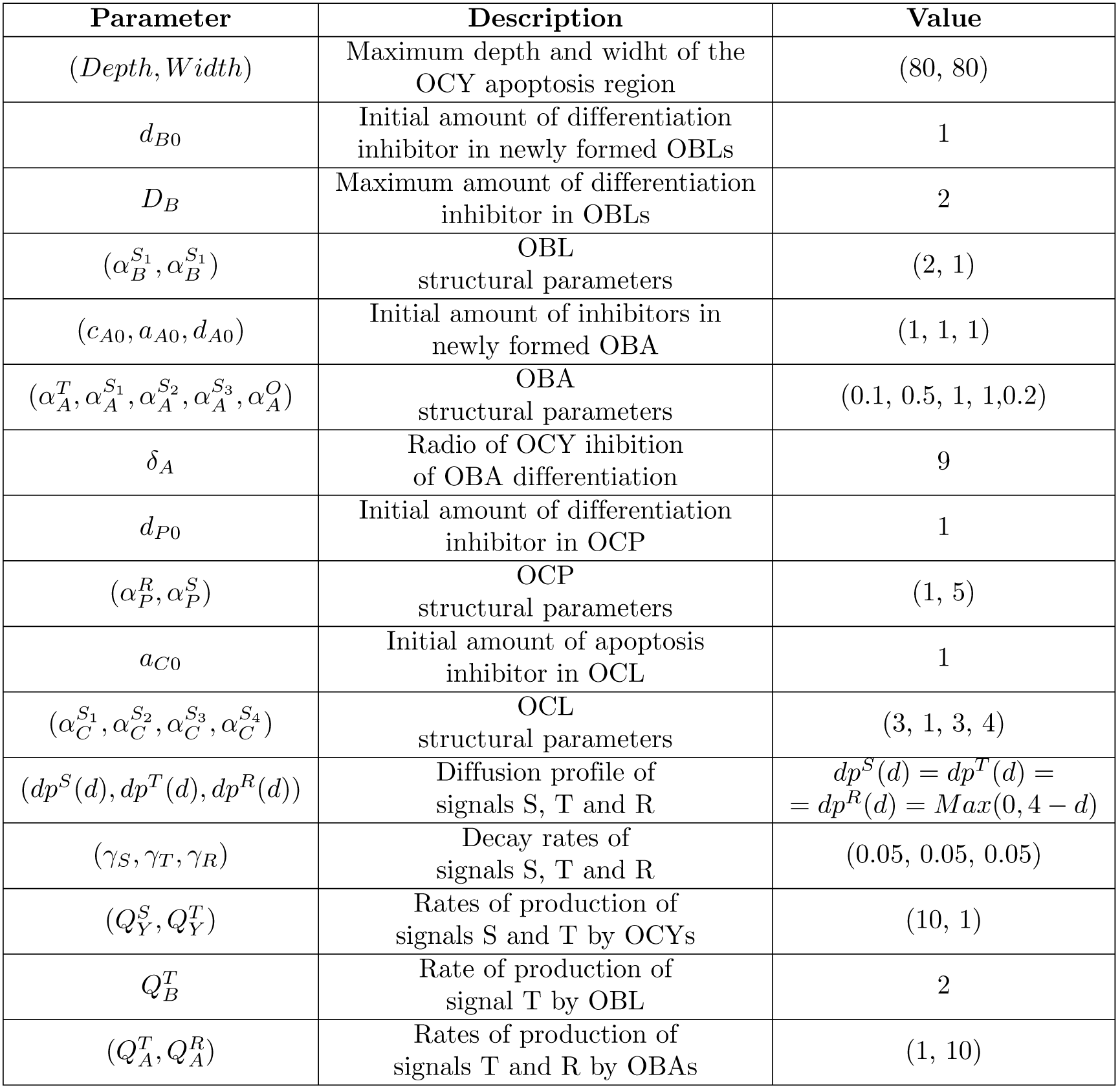
Values of the parameters used in the numerical simulations of the model.

## References

1. Graham JM, Ayati BP, Holstein SA, Martin JA. The role of osteocytes in targeted bone remodeling: a mathematical model. PLoS One. 2013;8(5):e63884. doi:10.1371/journal.pone.0063884.

2. Tate MLK, Adamson JR, Tami AE, Bauer TW. The osteocyte. The International Journal of Biochemistry & Cell Biology. 2004;36(1):1 – 8. doi:https://doi.org/10.1016/S1357-2725(03)00241-3.

3. Zaidi M. Skeletal remodeling in health and disease. Nature Medicine. 2007;13:791–801.

4. Safadi FF, Barbe MF, Abdelmagid SM, Rico MC, Aswad RA, Litvin J, et al. 1. In: Khurana JS, editor. Bone Structure, Development and Bone Biology. Totowa, NJ: Humana Press; 2009. p. 1–50. Available from: https://doi.org/10.1007/978-1-59745-347-9_1.

5. Jilka RL, Weinstein RS, Parfitt AM, Manolagas SC. Perspective: Quantifying Osteoblast and Osteocyte Apoptosis: Challenges and Rewards. Journal of Bone and Mineral Research. 2007;22(10):1492–1501. doi:10.1359/jbmr.070518.

6. Noble BS, Peet N, Stevens HY, Brabbs A, Mosley JR, Reilly GC, et al. Mechanical loading: biphasic osteocyte survival and targeting of osteoclasts for bone destruction in rat cortical bone. American Journal of Physiology-Cell Physiology. 2003;284(4):C934–C943. doi:10.1152/ajpcell.00234.2002.

7. Echeverri LF, Herrero MA, López JM, Oleaga G. Early stages of bone fracture healing: formation of a fibrin–collagen scaffold in the fracture hematoma. Bulletin of Mathematical Biology. 2015;77(1):156–183.

8. Hattner R, Epker BN, Frost HM. Suggested Sequential Mode of Control of Changes in Cell Behaviour in Adult Bone Remodelling. Nature. 1965;206(4983):489–490. doi:10.1038/206489a0.

9. Parfitt AM. Targeted and nontargeted bone remodeling: relationship to basic multicellular unit origination and progression. Bone. 2002;30(1):5–7. doi:10.1016/s8756-3282(01)00642-1.

10. Eriksen EF. Cellular mechanisms of bone remodeling. Reviews in Endocrine and Metabolic Disorders. 2010;11(4):219–227. doi:10.1007/s11154-010-9153-1.

11. Hauge EM, Qvesel D, Eriksen EF, Mosekilde L, Melsen F. Cancellous Bone Remodeling Occurs in Specialized Compartments Lined by Cells Expressing Osteoblastic Markers. Journal of Bone and Mineral Research. 2001;16(9):1575–1582. doi:10.1359/jbmr.2001.16.9.1575.

12. Kalfas IH. Principles of bone healing. Neurosurgical Focus. 2001;10(4):1–4.

13. Franz-Odendaal TA, Hall BK, Witten PE. Buried alive: How osteoblasts become osteocytes. Developmental Dynamics. 2005;235(1):176–190. doi:10.1002/dvdy.20603.

14. Sims NA, Martin TJ. Coupling the activities of bone formation and resorption: a multitude of signals within the basic multicellular unit. BoneKEy Reports. 2014;3. doi:10.1038/bonekey.2013.215.

15. Komarova SV, Smith RJ, Dixon SJ, Sims SM, Wahl LM. Mathematical model predicts a critical role for osteoclast autocrine regulation in the control of bone remodeling. Bone. 2003;33(2):206–215. doi:10.1016/s8756-3282(03)00157-1.

16. Ryser MD, Komarova SV, Nigam N. The Cellular Dynamics of Bone Remodeling: A Mathematical Model. SIAM Journal on Applied Mathematics. 2010;70(6):1899–1921. doi:10.1137/090746094.

17. Graham JM, Ayati BP, Holstein SA, Martin JA. The Role of Osteocytes in Targeted Bone Remodeling: A Mathematical Model. PLoS ONE. 2013;8(5):e63884. doi:10.1371/journal.pone.0063884.

18. Robling AG, Castillo AB, Turner CH. Biomechanical and molecular regulation of bone remodeling. Annual Review of Biomedical Engineering. 2006;8(1):455–498. doi:10.1146/annurev.bioeng.8.061505.095721.

19. Steeve KT, Marc P, Sandrine T, Dominique H, Yannick F. IL-6, RANKL, TNF-*α*/IL-1: interrelations in bone resorption pathophysiology. Cytokine & Growth Factor Reviews. 2004;15(1):49–60. doi:10.1016/j.cytogfr.2003.10.005.

20. Noble BS. The osteocyte lineage. Archives of Biochemistry and Biophysics. 2008;473(2):106 – 111. doi:https://doi.org/10.1016/j.abb.2008.04.009.

21. Chen G, Deng C, Li YP. TGF-*β* and BMP Signaling in Osteoblast Differentiation and Bone Formation. Int Journal of Biological Sciences. 2012;8(2):272–288.

22. Marsell R, Einhorn TA. The biology of fracture healing. Injury. 2011;42(6):551–555. doi:10.1016/j.injury.2011.03.031.

23. Wu M, Chen G, Li YP. TGF-*β* and BMP signaling in osteoblast, skeletal development and bone formation, homeostasis and disease. Bone Research. 2016;4(1). doi:10.1038/boneres.2016.9.

24. Dufour C, Holy X, Marie PJ. Transforming growth factor-*β* prevents osteoblast apoptosis induced by skeletal unloading via PI3K/Akt, Bcl-2, and phospho-Bad signaling. American Journal of Physiology-Endocrinology and Metabolism. 2008;294(4):E794–E801. doi:10.1152/ajpendo.00791.2007.

25. Heino TJ, Hentunen TA, Väänänen HK. Osteocytes inhibit osteoclastic bone resorption through transforming growth factor-*β*: Enhancement by estrogen. Journal of Cellular Biochemistry. 2002;85(1):185–197. doi:10.1002/jcb.10109.

26. Stein GS, Lian JB, Owen TA. Relationship of cell growth to the regulation of tissue-specific gene expression during osteoblast differentiation. The FASEB Journal. 1990;4(13):3111–3123. doi:10.1096/fasebj.4.13.2210157.

27. Bidwell JP, Yang J, Robling AG. Is HMGB1 an osteocyte alarmin? Journal of Cellular Biochemistry. 2008;103(6):1671–1680. doi:10.1002/jcb.21572.

28. Rahman MS, Akhtar N, Jamil HM, Banik RS, Asaduzzaman SM. TGF-*β*/BMP signaling and other molecular events: regulation of osteoblastogenesis and bone formation. Bone Research. 2015;3(1). doi:10.1038/boneres.2015.5.

29. Hock JM, Krishnan V, Onyia JE, Bidwell JP, Milas J, Stanislaus D. Osteoblast Apoptosis and Bone Turnover. Journal of Bone and Mineral Research. 2001;16(6):975–984. doi:10.1359/jbmr.2001.16.6.975.

30. Boyle WJ, Simonet WS, Lacey DL. Osteoclast differentiation and activation. Nature. 2003;423(6937):337–342.

31. Teitelbaum SL. Bone Resorption by Osteoclasts. Science. 2000;289(5484):1504–1508. doi:10.1126/science.289.5484.1504.

32. Mullen CA, Haugh MG, Schaffler MB, Majeska RJ, McNamara LM. Osteocyte differentiation is regulated by extracellular matrix stiffness and intercellular separation. Journal of the Mechanical Behavior of Biomedical Materials. 2013;28:183 – 194. doi:https://doi.org/10.1016/j.jmbbm.2013.06.013.

33. ABE E. Function of BMPs and BMP Antagonists in Adult Bone. Annals of the New York Academy of Sciences. 2006;1068(1):41–53. doi:10.1196/annals.1346.007.

34. Ishii T, Kikuta J, Kubo A, Ishii M. Control of osteoclast precursor migration: A novel point of control for osteoclastogenesis and bone homeostasis. IBMS BoneKEy. 2010;7(8):279–286. doi:10.1138/20100459.

35. Cao X. Targeting osteoclast-osteoblast communication. Nature Medicine. 2011;17(11):1344–1346. doi:10.1038/nm.2499.

36. Noble BS, Reeve J. Osteocyte function, osteocyte death and bone fracture resistance. Molecular and Cellular Endocrinology. 2000;159(1):7 – 13. doi:https://doi.org/10.1016/S0303-7207(99)00174-4.

37. You LD, Weinbaum S, Cowin SC, Schaffler MB. Ultrastructure of the osteocyte process and its pericellular matrix. The Anatomical Record. 2004;278A(2):505–513. doi:10.1002/ar.a.20050.

38. Christopher MJ, Link DC. Granulocyte Colony-Stimulating Factor Induces Osteoblast Apoptosis and Inhibits Osteoblast Differentiation. Journal of Bone and Mineral Research. 2008;23(11):1765–1774. doi:10.1359/jbmr.080612.

39. Reyes JP, Sims SM, Dixon SJ. P2 receptor expression, signaling and function in osteoclasts. Frontiers in bioscience (Scholar edition). 2011;3:1101—1118. doi:10.2741/214.

40. Karin M, Hunter T. Transcriptional control by protein phosphorylation: signal transmission from the cell surface to the nucleus. Current Biology. 1995;5(7):747 – 757. doi:https://doi.org/10.1016/S0960-9822(95)00151-5.

41. Jilka RL, Weinstein RS, Bellido T, Parfitt AM, Manolagas SC. Osteoblast Programmed Cell Death (Apoptosis): Modulation by Growth Factors and Cytokines. Journal of Bone and Mineral Research. 1998;13(5):793–802. doi:10.1359/jbmr.1998.13.5.793.

42. Arias CF, Herrero MA, Acosta FJ, Fernandez-Arias C. A mathematical model for a T cell fate decision algorithm during immune response. Journal of Theoretical Biology. 2014;349:109 – 120. doi:https://doi.org/10.1016/j.jtbi.2014.01.039.

43. Noman ASM, Koide N, Iftakhar-E-Khuda I, Dagvadorj J, Tumurkhuu G, Naiki Y, et al. Retinoblastoma protein-interacting zinc finger 1 (RIZ1) participates in RANKL-induced osteoclast formation via regulation of NFATc1 expression. Immunology Letters. 2010;131(2):166 – 169. doi:https://doi.org/10.1016/j.imlet.2010.04.006.

44. Tanaka S, Wakeyama H, Akiyama T, Takahashi K, Amano H, Nakayama KI, et al. Regulation of Osteoclast Apoptosis by Bcl-2 Family Protein Bim and Caspase-3. In: Choi Y, editor. Osteoimmunology. Boston, MA: Springer US; 2010. p. 111–116.

45. Haxaire C, Haӱ E, Geoffroy V. Runx2 Controls Bone Resorption through the Down-Regulation of the Wnt Pathway in Osteoblasts. The American Journal of Pathology. 2016;186(6):1598 – 1609. doi:https://doi.org/10.1016/j.ajpath.2016.01.016.

46. Chau JFL, Leong WF, Li B. Signaling pathways governing osteoblast proliferation, differentiation and function. Histology and histopathology. 2009;24(12):1593–1606.

47. Marotti G, Ferretti M, Muglia MA, Palumbo C, Palazzini S. A quantitative evaluation of osteoblast-osteocyte relationships on growing endosteal surface of rabbit tibiae. Bone. 1992;13(5):363 – 368. doi:https://doi.org/10.1016/8756-3282(92)90452-3.

48. Nelson DL, Lehninger AL, Cox MM. Lehninger Principles of Biochemistry. Macmillan; 2008.

49. Ebenhöh O, Heinrich R. Evolutionary optimization of metabolic pathways. Theoretical reconstruction of the stoichiometry of ATP and NADH producing systems. Bulletin of Mathematical Biology. 2001;63(1):21–55.

50. Hemker HC, Kerdelo S, Kremers RMW. Is there value in kinetic modeling of thrombin generation? No (unless…). Journal of Thrombosis and Haemostasis. 2012;10(8):1470–1477. doi:10.1111/j.1538-7836.2012.04802.x.

51. Ginzburg LR, Jensen CXJ. Rules of thumb for judging ecological theories. Trends in Ecology & Evolution. 2004;19(3):121 – 126. doi:https://doi.org/10.1016/j.tree.2003.11.004.

52. Bezanson J, Edelman A, Karpinski S, Shah VB. Julia: A Fresh Approach to Numerical Computing. SIAM Review. 2017;59(1):65–98. doi:10.1137/141000671.

